# The Infraslow Fluctuation of Sigma Power During Sleep in Young Individuals with Schizophrenia

**DOI:** 10.1101/2025.04.23.650209

**Authors:** Maria E. Dimitriades, Eve Schumacher, Janani Arudchelvam, Sara Fattinger, Salome Kurth, Fiona Pugin, Flavia Wehrle, Valeria Jaramillo, Carina Volk, Sven Leach, Ashura Buckley, David Driver, Andjela Markovic, Judith Rapoport, Leila Tarokh, Reto Huber, Miriam Gerstenberg

**Author notes:** (first author).

## Abstract

A reduction in sleep spindles, a major electrophysiological characteristic of Non-Rapid Eye Movement sleep, has been suggested as a potential biomarker of schizophrenia. While research has primarily focused on the spindle quantity, recent studies have begun to explore their temporal dynamics throughout the night. In healthy individuals, sleep spindles fluctuate on an infraslow ∼50-second timescale, alternating between phases of high and low spindle activity. This fluctuation is referred to as the infraslow fluctuation of sigma power (ISFS), which is modulated by noradrenergic activity from the locus coeruleus and linked to the organization of arousal and memory reactivation processes during sleep. Given the known deficit in sleep spindles, dysregulation of noradrenergic activity, and impairments in sleep maintenance and memory in schizophrenia, this study investigates the ISFS in sleep electroencephalography data from individuals with either Childhood-Onset Schizophrenia (COS; N = 17) or Early-Onset Schizophrenia (EOS; N = 11), aged 9 to 21 years, alongside age- and sex-matched healthy controls (N = 56). The presence and strength of the ISFS were reduced in both COS and EOS groups compared to controls, particularly in central-parietal electrodes. No significant differences in these features of the ISFS were found between the two clinical groups, despite group differences in sleep spindle density and clinical characteristics. These findings suggest that the ISFS is observable but reduced in young patients with schizophrenia and support the notion that the timing of sleep spindles may inform pathomechanistic models of the disorder, as well as future diagnostic approaches and interventions.

## Introduction

A reduction in sleep spindles—brief oscillatory bursts characteristic of Non-Rapid Eye Movement (NREM) sleep—has been consistently reported in schizophrenia (Castelnovo et al., 2018; Ferrarelli, 2024; Ferrarelli et al., 2007; Manoach et al., 2016). Over the past two decades, this spindle deficit has been attributed to the impairment of glutamatergic thalamocortical circuits in schizophrenia (Benson, 2006; Manoach et al., 2016; Vukadinovic, 2011; Zhang et al., 2020). The reduction in sleep spindles is observed in adult-onset schizophrenia when compared to both healthy and clinical controls and appears independent of medication status (Castelnovo et al., 2018). Sleep spindle deficits have been observed even in young individuals with schizophrenia, including those with Childhood-Onset Schizophrenia (COS; diagnosis before age 13) and Early-Onset Schizophrenia (EOS; diagnosis between ages 13 and 18) (Dimitriades et al., 2023; Gerstenberg et al., 2020; Markovic et al., 2020). Although rare in childhood, schizophrenia incidence rises sharply during adolescence (Solmi et al., 2021). Studying young individuals with schizophrenia is especially important, as early identification of biomarkers could help overcome diagnostic delays common in these individuals, enabling earlier intervention and improving long-term outcomes (Schimmelmann et al., 2007, 2008). Moreover, adolescence represents a particularly vulnerable period, when ongoing brain maturation and network specialization may increase sensitivity to pathological disruptions—while also offering a potential window for more effective therapeutic intervention (Gogtay et al., 2011; Insel, 2010; Pantelis et al., 2005).

Recently, growing interest has shifted beyond the number of sleep spindles to their precise timing (Champetier et al., 2023; Dimitriades et al., 2024; Lázár et al., 2019; Lecci et al., 2017; Osorio-Forero et al., 2021). Rather than occurring evenly throughout NREM sleep, sleep spindles fluctuate over an approximately 50-second timescale, alternating between phases of high and low spindle activity (Lecci et al., 2017; Watson, 2018). This variation in the timing of sleep spindle activity has been observed in rodents as well as humans, spanning from childhood to late adulthood, and is referred to as the infraslow fluctuation of sigma power (ISFS) (Champetier et al., 2023; Dimitriades et al., 2024; Lecci et al., 2017). In rodents, the ISFS has been found to be driven by locus coeruleus noradrenergic activity, a key modulator of arousal in the brain (Hayat et al., 2020; Osorio-Forero et al., 2021). Interestingly, arousals during sleep—as well as markers of memory reactivation—occur at specific phases of the ISFS in both rodents and humans (Dimitriades et al., 2024; Lecci et al., 2017), suggesting that there may be optimal brain states for efficient arousal regulation and memory processing during sleep. These findings have sparked interest in examining the ISFS in clinical populations, where such processes are often disrupted.

Young individuals with schizophrenia have not been well studied with respect to arousals during sleep or memory reactivation, whereas adults with chronic schizophrenia exhibit more frequent arousals (Bagautdinova et al., 2023) and reduced or absent overnight improvements in procedural learning tasks, when compared to healthy controls (Genzel et al., 2015; Manoach et al., 2004; Seeck-Hirschner et al., 2010). Moreover, while predominant hypotheses on the pathomechanisms of schizophrenia focus on disruptions in glutamatergic and dopaminergic signaling (Howes et al., 2015; Kitzinger & Arnold, 1949; Lieberman et al., 1987), there is also evidence pointing to dysfunction in the locus coeruleus-noradrenaline system (Mäki-Marttunen et al., 2020; Maletic et al., 2017; Yamamoto & Hornykiewicz, 2004).

Given the known deficits in sleep spindles, alongside dysregulated noradrenergic activity and increased arousals and impairments in memory, schizophrenia presents a compelling case for investigating potential alterations in the ISFS. To explore this, electroencephalographic (EEG) sleep data were gathered from two independent clinical samples of young individuals with schizophrenia (one group with COS, one with EOS). Features of the ISFS, specifically its presence and strength, were compared between patients and healthy controls, as well as between the two patient groups, and potential associations between features of the ISFS, sleep spindle density, and clinical characteristics were examined.

## Materials and Methods

### Participants

This project pooled sleep, demographic, and clinical data from two patient groups diagnosed with schizophrenia. One sample was diagnosed with Childhood-Onset Schizophrenia (COS; N = 17), implying that the age of onset occurred before 13 years. These data were recorded as a follow-up after a few years of illness (mean ± standard deviation (SD) age at assessment: 16.00 ± 3.64 years; sex: 70.59% female, 29.41% male; mean duration of illness: 6.73 years). The other sample, diagnosed with Early-Onset Schizophrenia (EOS; N = 11), had an age of onset between 13 and 18 years and was recorded shortly after diagnosis (age at assessment: 16.38 ± 1.42 years; sex: 36.36% female, 63.64% male; mean duration of illness: 1.05 years). All individuals were aged between 9 and 21 years at the time of the sleep EEG.

The COS group was recruited at the National Institute of Mental Health in the United States between 2014 and 2017 and the EOS group at the Department of Child and Adolescent Psychiatry of the Psychiatric University Hospital Zurich in Switzerland between 2012 and 2016. Screening procedures and characteristics of these patient groups have been previously published (Dimitriades et al., 2023; Gerstenberg et al., 2020; Markovic et al., 2020). This research project was approved by Ethics Committee of the Canton of Zurich and Bern, Switzerland, BASEC 2021-00215. Written informed consent to participate in this study was provided by the participants’ legal guardian/next of kin. All study procedures were performed according to the Declaration of Helsinki. Age of illness onset (in years), duration of illness (in years), antipsychotic medication (measured as chlorpromazine equivalent), and positive and negative symptomatology were available for every individual with schizophrenia, either according to the Scale for the Assessment of Positive Symptoms (SAPS) and Scale for the Assessment of Negative Symptoms (SANS) for the COS group or to the Positive and Negative Symptoms Scale (PANSS) for the EOS group (Andreasen, 1986, 1989; Kay et al., 1987).

Every individual with COS and EOS was age- and sex-matched to two healthy controls (HC). This resulted in 34 individuals in the HC group for the COS group (HC-COS; 16.10 ± 3.86 years; 70.59% female, 29.41% male) and 22 individuals in the HC group for the EOS group (HC-EOS; 16.46 ± 1.62 years; 36.36% female, 63.64% male). Sleep and demographic data for all controls were collected between 2008 and 2021 in the sleep laboratory of the University Children’s Hospital Zurich in Switzerland (Groch et al., 2016; Kurth et al., 2010; Leach et al., 2024; Pugin et al., 2015; Volk et al., 2018, 2019; Wehrle et al., 2017). Sex was recorded as a binary variable (female or male).

### Electroencephalogram recordings and preprocessing

EEG recording systems and sleep-specific preprocessing varied between the COS group (Markovic et al., 2020) and the EOS group and all HCs (Gerstenberg et al., 2020; Groch et al., 2016; Kurth et al., 2010; Leach et al., 2024; Pugin et al., 2015; Volk et al., 2018, 2019; Wehrle et al., 2017). All-night EEG were recorded from patients with COS using a 21-channel 10-20 system (TWin or Nihon Kohden; sampling rate: 200 Hz) and from patients with EOS and all HCs using a 128-channel system (Electrical Geodesic Sensory Net; sampling rate: 500 Hz). There were 21 channels in common between all individuals, as previously reported (Dimitriades et al., 2023).

Sleep stages were scored according to standard criteria by two sleep experts (Berry et al., 2020). Each epoch was assigned a standard sleep stage, with epochs lasting 30 seconds for the COS group and 20 seconds for the EOS group and all HCs. Semi-automatic artifact rejection was applied based on power in the delta and beta frequency bands, following previously established methods (Huber et al., 2000; Leach et al., 2023). Only artifact-free epochs across all channels were included in further analyses. Channels with poor signal quality were interpolated from neighboring channels. Further, all EEG data were low-pass filtered (−3 dB cut-off: 39.25 Hz; Hamming Window Finite Impulse Response (FIR) filter; filter order: 184; *pop_eegfiltnew* from EEGLAB) (Delorme & Makeig, 2004), downsampled (200 Hz; for the EOS group and all HC; *pop_resample* from EEGLAB), and high-pass filtered (−3 dB cut-off: 0.27 Hz; Kaiser Window FIR filter; filter order: 2390; custom implementation in MATLAB). The data were re-referenced to the mean of the mastoids.

### Analyses of the infraslow fluctuation of sigma power and sleep spindles

Uninterrupted NREM Stage N2 (N2) sleep, free from artifacts or transitions to other vigilance states, was isolated. This procedure followed Dimitriades et al. (2024), with the exception that N2 sleep bouts were required to last at least 300 seconds (instead of 280), to account for differences in sleep staging epoch lengths between groups (20 versus 30 seconds). N2 sleep bout parameters were compared between patients and controls. These parameters include: the number of bouts, the mean bout length (in seconds), the mean relative location of the bouts (the mean across all bout midpoints divided by the total length of the recording; in percent), the total duration of the bouts (in minutes), and the proportion of all N2 (the duration of the N2 bout data is divided by the total duration of N2 sleep and multiplied by 100; in percent). The subsequently explained ISFS and sleep spindle analyses were performed on the aforementioned N2 sleep bouts.

The ISFS was analyzed as previously described (Dimitriades et al., 2024). In brief, the sigma power was extracted over time from N2 sleep bouts using a Gabor-Morlet wavelet (4 cycles, 0.2 Hz resolution) across a broad sigma range (10-16 Hz) to capture frontal and central sleep spindle activity (Fernandez & Lüthi, 2020; Kwon et al., 2023). The sigma power over time was then centered around zero by subtracting the mean, and the Fast Fourier Transform (FFT) was applied. To account for varying N2 sleep bout lengths (≥300 seconds), the spectral power was adjusted by dividing the FFT output by the given bout duration. Spectral power was averaged across all N2 bouts. To account for potential intra- and interindividual differences, the average spectral power was normalized by dividing spectral power in each frequency bin by the mean spectral power across all frequency bins (0-0.1 Hz). A horizontal baseline correction was further applied by subtracting the mean power from 0.06 to 0.1 Hz. The frequency range for the ISFS peak detection was set between 0.01 and 0.04 Hz, where the infraslow spectral peak typically occurs (Lecci et al., 2017). This procedure was performed on every channel and participant separately.

Subsequently, two features of the ISFS, the presence and the strength, were calculated for each channel and participant. To assess the presence of the ISFS, two criteria were applied: 1) a Gaussian function was attempted to be fit on the spectral power and 2) if a Gaussian was fit, a threshold (1.5 times the SD of the Gaussian) was defined that the peak power must surpass. If both criteria were met, the presence of the ISFS was confirmed in that given channel and participant (Dimitriades et al., 2024). To assess the strength of the ISFS, the area under the curve of the spectral power was calculated using the MATLAB function *trapz*, which approximates the integral of the spectral power. Spectral powers between 0.01 and 0.04 Hz were considered (Lecci et al., 2017; Osorio-Forero et al., 2021).

Sleep spindles were detected using an amplitude-based thresholding approach (Ferrarelli et al., 2007). N2 sleep bout data were first extracted and processed with a band-pass filter (−3 dB cut-offs: 9.96 and 17.5 Hz; Chebyshev Type II IIR filter; filter order: 6; using the MATLAB function *filtfilt*). Sleep spindles were further identified as periods when the signal amplitude exceeded the upper threshold and continued until it dropped below the lower threshold. The upper threshold corresponded to five times the channel-specific mean amplitude and the lower threshold to two times the channel-specific mean amplitude. Only events lasting at least 0.3 seconds were considered sleep spindles (Warby et al., 2014). Sleep spindle density was calculated as the number of detected sleep spindles divided by the total time of the N2 sleep bout data in minutes.

### Statistical analyses

All statistical analyses were performed in MATLAB R2022a. Data normality was evaluated using quantile-quantile plots and Shapiro-Wilk tests. Group differences in age and sleep structure parameters were assessed using Mann-Whitney U tests for non-parametric data, with effect sizes estimated via Cliff’s delta, while sex differences were assessed using Fisher’s exact test. The percent of electrodes showing the presence of the ISFS was compared between patient and control groups using an unpaired t-test, while the presence of the ISFS at each electrode was compared between patient and control groups using a z-test for proportions. Differences in the strength of the ISFS and sleep spindle density were analyzed with unpaired t-tests. To account for multiple comparisons across 19 channels, we defined a minimum cluster size based on an alpha rate of 5%, ensuring that isolated significant electrodes due to chance were excluded. Therefore, only clusters of at least two adjacent significant electrodes were considered meaningful. Pearson correlations were used to examine relationships between symptomatology and the strength of the ISFS in patients (extracted from a cluster of central-parietal channels: C3, Cz, C4, P3, Pz, P4) and sleep spindle density and the strength of the ISFS in all groups (compared in one central channel: C4). If not otherwise specified, values are expressed as the mean ± SD.

## Results

### Demographic and sleep structure parameters

Demographic and all-night sleep structure parameters have been previously published (Dimitriades et al., 2023). Unlike in the previous study, each patient in this project was matched with two healthy controls instead of one to enhance statistical robustness. Age and sex were not significantly different between the patient groups and their respective healthy control groups (**Table 1**). All-night sleep structure parameters were not significantly different for the EOS group and the HC-EOS group. The percent of time spent in NREM sleep stage N2 and N3 and rapid eye movement (REM) were not significantly different between the COS group and the HC-COS group. The COS group had a significantly smaller percentage of wake after sleep onset (COS mean ± SD: 1.44 ± 1.47%; HC-COS: 6.06 ± 7.54%; Cliff’s Delta = -0.65, p < 0.001) and N1 sleep (COS: 3.59 ± 2.74%; HC-COS: 6.81 ± 4.31%; Cliff’s Delta = -0.54, p < 0.01) compared to the HC-COS group. The COS group also spent significantly more time asleep when compared to the HC-COS group (COS: 542.62 ± 157.73 minutes; HC-COS: 452.93 ± 51.83 minutes; Cliff’s Delta = 0.65, p < 0.001). These differences have been previously reported (Markovic et al., 2020).

**Table 1:** Demographic and sleep structure parameters. COS: Childhood-Onset Schizophrenia; EOS: Early-Onset Schizophrenia; HC-COS: Healthy Controls for COS; HC-EOS: Healthy Controls for EOS; SD: Standard Deviation; WASO: Wake After Sleep Onset; Non-Rapid Eye Movement Sleep Stages (N1, N2, N3); REM: Rapid Eye Movement; min: minutes; s: seconds; %: percent. Sex is reported as a binary variable (female or male), with the number of females indicated in the table. Bout: N2 sleep data lasting at least 300 seconds without transitions to other vigilance states or artifacts. Mean Relative Location: Mean of all bout midpoints divided by the total recording length, expressed as a percentage. Proportion of all N2: Total duration of N2 bout data relative to total N2 sleep duration. Statistically significant p-values are shown in bold.

| Parameter | Patient Group (N) | Mean (SD) | Healthy Controls (N) | Mean (SD) | Cliff's Delta | p |
| --- | --- | --- | --- | --- | --- | --- |
| <u>Demographics</u> |  |  |  |  |  |  |
| Age (years) | COS (17) | 16.00 (3.64) | HC-COS (34) | 16.10 (3.86) | 0.02 | 0.90 |
|  | EOS (11) | 16.38 (1.42) | HC-EOS (22) | 16.46 (1.62) | 0.00 | 1.00 |
| Sex (N of females) | COS (17) | 12 | HC-COS (34) | 24 | 0.00 | 1.00 |
|  | EOS (11) | 4 | HC-EOS (22) | 8 | 0.00 | 1.00 |
| <u>All-Night Sleep</u> |  |  |  |  |  |  |
| WASO (%) | COS (17) | 1.44 (1.47) | HC-COS (34) | 6.06 (7.54) | -0.65 | **<0.001** |
|  | EOS (11) | 5.31 (4.54) | HC-EOS (22) | 5.52 (6.09) | -0.03 | 0.89 |
| N1 (%) | COS (17) | 3.59 (2.74) | HC-COS (34) | 6.81 (4.31) | -0.54 | **<0.01** |
|  | EOS (11) | 6.71 (3.93) | HC-EOS (22) | 7.15 (5.00) | 0.00 | 1.00 |
| N2 (%) | COS (17) | 48.50 (19.93) | HC-COS (34) | 50.19 (7.05) | 0.02 | 0.91 |
|  | EOS (11) | 48.05 (10.23) | HC-EOS (22) | 52.22 (4.50) | -0.21 | 0.33 |
| N3 (%) | COS (17) | 29.64 (24.52) | HC-COS (34) | 22.50 (8.43) | 0.06 | 0.74 |
|  | EOS (11) | 26.07 (13.01) | HC-EOS (22) | 20.63 (6.02) | 0.20 | 0.37 |
| REM (%) | COS (17) | 18.27 (9.78) | HC-COS (34) | 20.51 (5.18) | -0.20 | 0.26 |
|  | EOS (11) | 19.16 (7.70) | HC-EOS (22) | 20.00 (5.33) | 0.04 | 0.86 |
| Total Sleep Time (min) | COS (17) | 542.62 (167.73) | HC-COS (34) | 452.93 (51.83) | 0.65 | **<0.001** |
|  | EOS (11) | 437.85 (123.67) | HC-EOS (22) | 417.26 (63.55) | 0.37 | 0.09 |
| <u>Bouts of N2 Sleep</u> |  |  |  |  |  |  |
| Bout Number | COS (17) | 13.71 (10.09) | HC-COS (34) | 12.71 (3.24) | 0.02 | 0.92 |
|  | EOS (11) | 13.64 (5.82) | HC-EOS (22) | 13.00 (3.27) | 0.17 | 0.45 |
| Mean Bout Length (s) | COS (17) | 450.58 (86.42) | HC-COS (34) | 588.37 (121.86) | -0.68 | **<0.0001** |
|  | EOS (11) | 531.96 (96.48) | HC-EOS (22) | 576.95 (104.70) | -0.24 | 0.28 |
| Mean Relative Location (%) | COS (17) | 52.23 (13.34) | HC-COS (34) | 54.16 (7.16) | -0.08 | 0.67 |
|  | EOS (11) | 54.10 (7.50) | HC-EOS (22) | 51.09 (7.05) | 0.25 | 0.26 |
| Total Duration of Bouts (min) | COS (17) | 110.35 (91.19) | HC-COS (34) | 127.21 (47.00) | -0.20 | 0.26 |
|  | EOS (11) | 123.06 (55.57) | HC-EOS (22) | 123.98 (34.86) | 0.14 | 0.54 |
| Proportion of all N2 (%) | COS (17) | 38.23 (23.40) | HC-COS (34) | 55.07 (15.71) | -0.41 | **0.02** |
|  | EOS (11) | 56.01 (15.43) | HC-EOS (22) | 56.68 (11.29) | -0.02 | 0.95 |

**Table 2:** Clinical characteristics between groups with schizophrenia. COS: Childhood-Onset Schizophrenia; EOS: Early-Onset Schizophrenia. Age of onset and duration of illness are measured in years and antipsychotic medication is measured as the chlorpromazine equivalent. Statistically significant p-values are shown in bold.

| Parameter | EOS (11):<br>Mean (SD) | COS (17):<br>Mean (SD) | Cliff's Delta | p |
| --- | --- | --- | --- | --- |
| Age of Onset (years) | 15.33 (1.45) | 9.27 (1.79) | 1.00 | **<0.0001** |
| Duration (years) | 1.05 (0.92) | 6.73 (3.63) | -0.94 | **<0.0001** |
| Antipsychotic Medication (mg) | 172.50 (158.43) | 667.65 (372.05) | -0.78 | **<0.001** |

N2 bout sleep parameters—including the number of bouts, mean bout length, mean relative bout location, total bout duration, and the proportion of time spent in N2 bout sleep relative to total N2 sleep—were compared between schizophrenia and control groups (**Table 1**). The EOS and HC-EOS groups were not significantly different for all N2 bout sleep parameters. The COS and HC-COS groups were not significantly different in terms of the number of bouts, mean relative bout location, and the total bout duration. In contrast, the COS group had a significantly shorter mean bout length (COS: 450.58 ± 86.42 seconds; HC-COS: 588.37 ± 121.86 seconds; Cliff’s Delta = -0.68, p < 0.0001) and they spent a smaller proportion of their total N2 sleep across the night in N2 sleep bouts (COS: 38.23 ± 23.40%; HC-COS: 55.07 ± 15.71%; Cliff’s Delta = -0.41, p < 0.05) when compared to the HC-COS group.

### Presence of the infraslow fluctuation of sigma power is reduced in patients with schizophrenia compared to healthy controls

In a first step, the presence of the ISFS was quantified for each electrode and participant. As expected, healthy controls showed the presence of the ISFS in most electrodes (percent of electrodes with the ISFS: HC-COS: 73.99 ± 27.64%; HC-EOS: 82.54 ± 19.79%). In contrast, the percent of electrodes with the presence of the ISFS was significantly lower in the COS (54.49 ± 28.76%; t(49) = 2.34, p < 0.05) and EOS (59.33 ± 34.12%; t(31) = 2.48, p < 0.05) groups when compared to their respective control group.

To further investigate the spatial characteristics of the presence of the ISFS, its topographical distribution across the scalp was examined. In both control groups, the presence of the ISFS was maximal over central-parietal regions, whereas no clear topographical pattern emerged in either schizophrenia group. The presence of the ISFS was significantly reduced in the COS group compared to the HC-COS group and in the EOS group compared to the HC-EOS group, primarily in central-parietal regions (**Figure 1A**).

**Figure 1:**
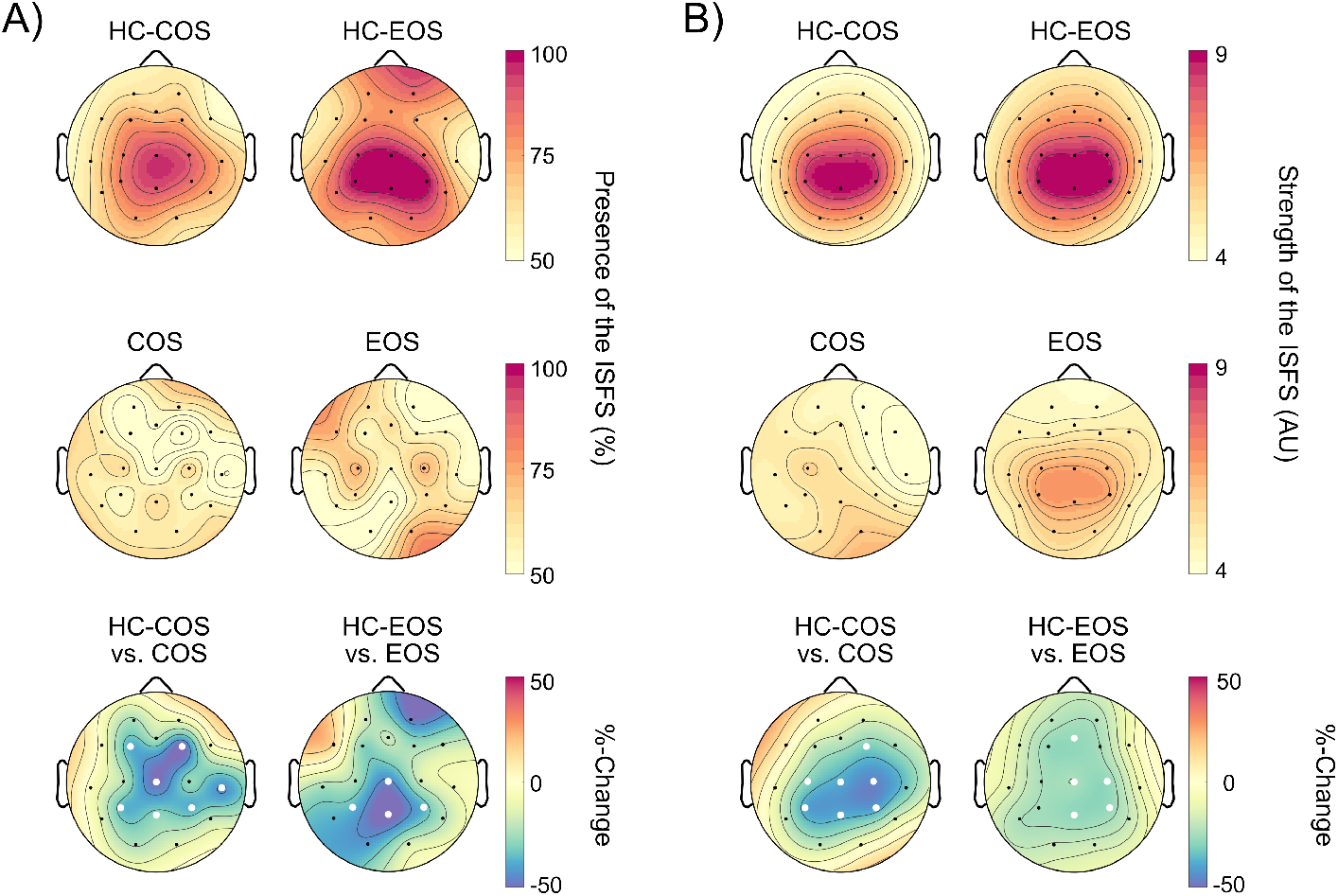
Individuals with schizophrenia show a reduction in features of the infraslow fluctuation of sigma power Topographical distribution of features of the infraslow fluctuation of sigma power (ISFS) in patient groups (middle row: COS: Childhood-Onset Schizophrenia; EOS: Early-Onset Schizophrenia) and their respective healthy control groups (top row: HC-COS: Healthy Controls for COS; HC-EOS: Healthy Controls for EOS) are presented. A) Presence of the ISFS is expressed as the percentage of individuals in each group exhibiting the ISFS at a given electrode. For example, at electrode Cz, 100% of the HC-EOS group (22 of 22 individuals) show the ISFS. Group differences in the presence of the ISFS were assessed using a z-test for proportions (electrodes showing significant group differences are marked with a white dot) and presented as the percent change between groups (bottom left). B) Strength of the ISFS was compared using an unpaired t-test (electrodes showing significant group differences are marked with a white dot) and is presented as the percent change between groups (bottom right).

### Strength of the infraslow fluctuation of sigma power is reduced in patients with schizophrenia compared to healthy controls

To overcome the weakness of analyzing a binary variable (i.e., the presence of the ISFS), in the next step, the strength of the ISFS was quantified for each electrode and participant. In both control groups, the topography of the strength of the ISFS closely aligns with the topography of the presence of the ISFS, with maximal values over central-parietal regions (specifically C3, Cz, C4, P3, Pz, P4). The EOS group exhibited a similar topography to controls but with a lower magnitude, whereas the COS group showed no discernible topographical pattern. The strength of the ISFS was significantly reduced in the COS group compared to the HC-COS group and in the EOS group compared to the HC-EOS group, primarily in central-parietal regions (**Figure 1B**).

To assess the consistency of these findings across individuals, the strength of the ISFS was plotted for each patient with schizophrenia alongside their age- and sex-matched healthy controls (**Figure 2**). Most healthy controls exhibited local maxima in the previously identified central-parietal cluster, a pattern that was less pronounced in the schizophrenia groups.

**Figure 2:**
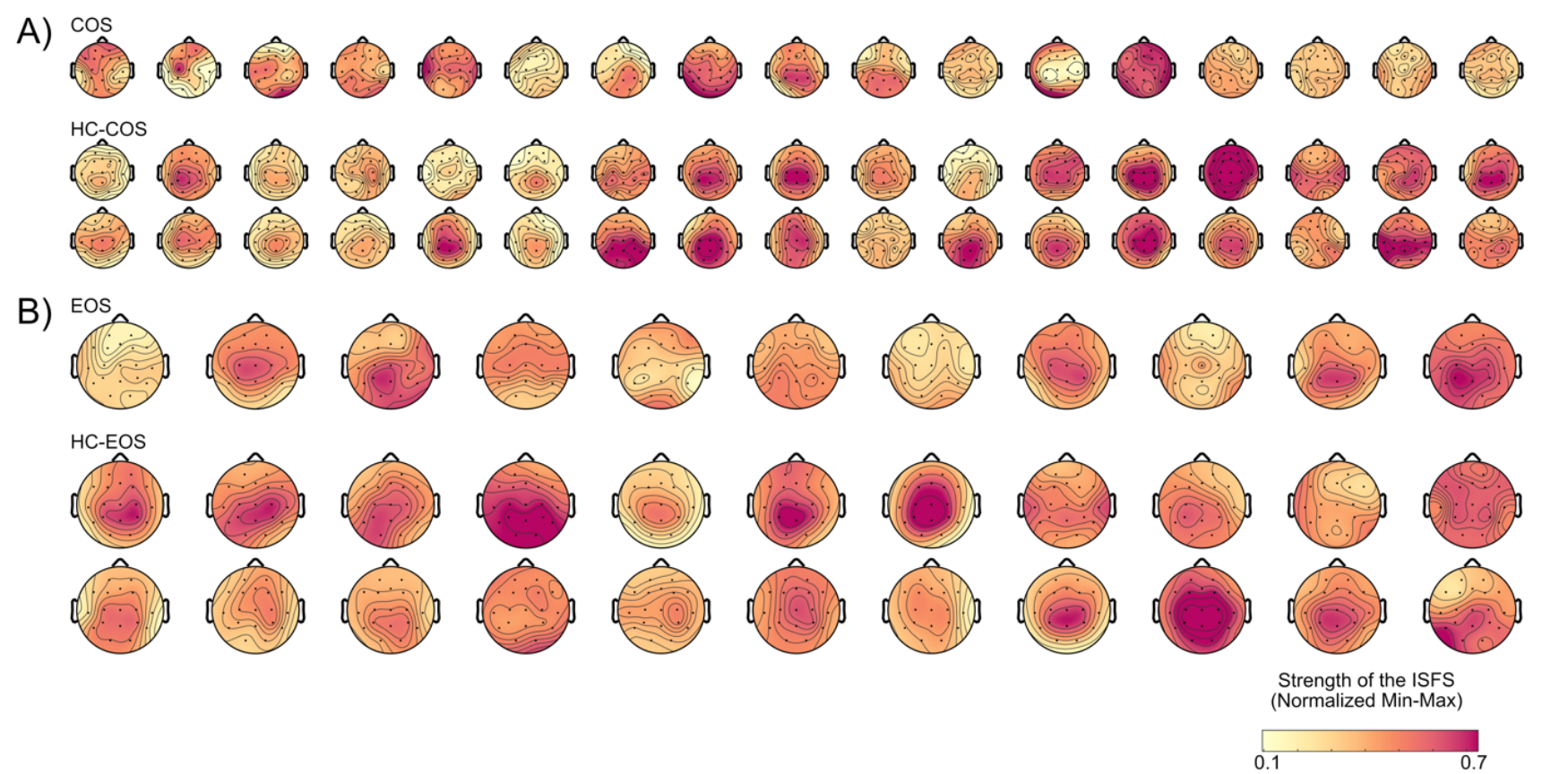
Individual topoplots of the strength of the infraslow fluctuation in sigma power for all patients with schizophrenia and healthy controls Topographical distribution of the strength of the infraslow fluctuation of sigma power (ISFS) is shown A) for every individual with Childhood-Onset Schizophrenia (COS; top) and their two age- and sex-matched healthy controls (HC-COS; bottom) and B) for every individual with Early-Onset Schizophrenia (EOS; top) and their two age- and sex-matched controls (HC-EOS; bottom). The strength of the ISFS values were normalized using Min-Max scaling, rescaling each value to a range of 0 to 1 based on the global minimum and maximum values across groups. Individuals are plotted by age (youngest on the left, oldest on the right).

### Features of the infraslow fluctuation of sigma power are not significantly different between the schizophrenia groups

To assess potential differences in the presence and strength of the ISFS between the patient groups, individuals of the COS and EOS groups were one-to-one matched. Since the age range differed between groups (EOS age range: 13.50 to 17.92 years; COS age range: 9.00 to 21.00 years), the youngest three and oldest three COS participants were excluded (COS: 16.00 ± 2.37 years); all individuals from the EOS group were included (EOS: 16.38 ± 1.42 years). This resulted in similar age distribution while sex-matching was not applied.

This approach showed that the presence of the ISFS did not differ significantly between the COS and EOS groups at any electrode (p > 0.10). Similarly, the strength of the ISFS did not significantly differ between the COS and EOS groups at any electrode (p > 0.23).

### No association between the strength of the infraslow fluctuation of sigma power and sleep spindle density

To examine the potential relationship between the strength of the ISFS and sleep spindle density (number of sleep spindles per minute), the extent of the sleep spindle deficit was assessed in each patient group and correlation analyses between the strength of the ISFS and sleep spindle density were performed.

As previously shown (Dimitriades et al., 2023; Markovic et al., 2020), the COS group demonstrated an almost global reduction in sleep spindle density compared to the HC-COS group (**Figure 3**). In contrast, the EOS group showed no significant difference in sleep spindle density when compared to the HC-EOS group. Also, in the one-to-one matching between the COS and EOS groups, sleep spindle density was significantly lower in the COS group than in the EOS group across all frontal, central, and parietal electrodes (p < 0.05), while no significant differences were observed in temporal and occipital electrodes (p > 0.62).

**Figure 3:**
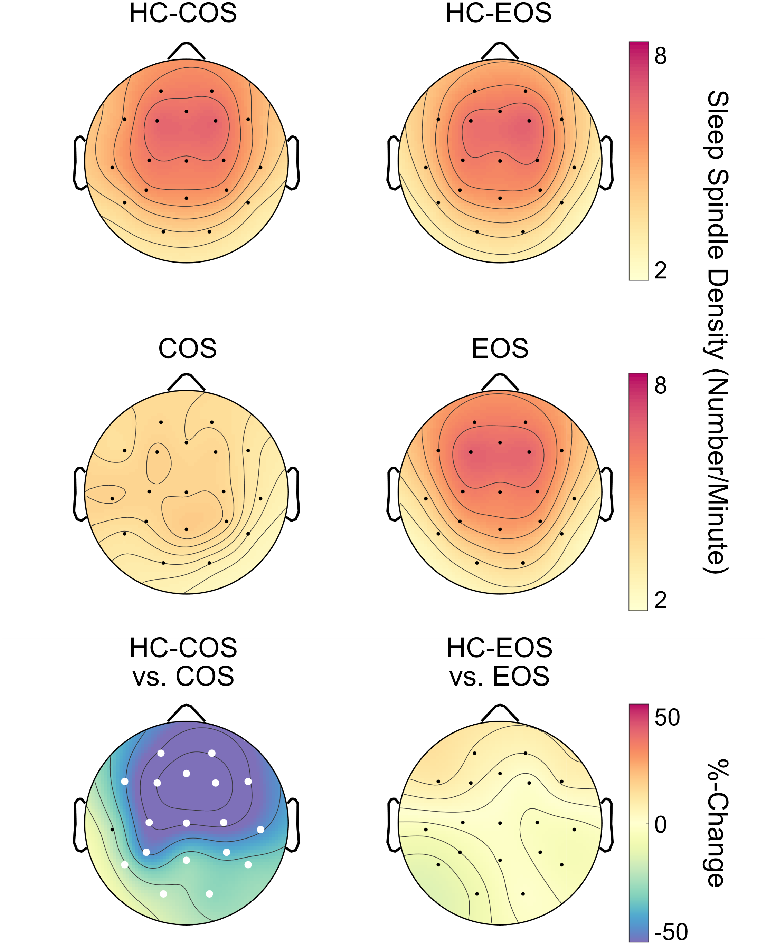
Sleep spindle density in non-rapid eye movement stage N2 sleep bout data Topographical distribution of sleep spindle density (the number of sleep spindles per minute) in patient groups (middle row: COS: Childhood-Onset Schizophrenia; EOS: Early-Onset Schizophrenia) and their respective healthy control groups (top row: HC-COS: Healthy Controls for COS; HC-EOS: Healthy Controls for EOS). Sleep spindle density was compared using an unpaired t-test (electrodes showing significant group differences are marked with a white dot) and is presented as the percent change between groups (bottom row).

Additionally, there was no significant correlation between the strength of the ISFS and sleep spindle density at a central electrode (C4) for the COS group (rho = -0.15, p = 0.56) and the pooled healthy controls (rho = 0.07, p = 0.59); the EOS group showed a positive trend-level association (rho = 0.53, p = 0.09).

### No association between the strength of the infraslow fluctuation of sigma power and clinical characteristics

Comparison of clinical characteristics between the COS and EOS groups at the time of the sleep EEG showed that the COS group had a longer duration of illness (COS: 6.73 ± 3.63 years; EOS: 1.05 ± 0.92 years; Cliff’s Delta = -0.94, p < 0.0001) and higher amounts of the chlorpromazine equivalent (COS: 667.65 ± 372.05 milligrams; EOS: 172.50 ± 158.43 milligrams; Cliff’s Delta = -0.78, p < 0.001). As expected, given the definitions of COS and EOS, the age at onset was significantly lower in the COS group when compared to the EOS group (COS: 9.27 ± 1.79 years; EOS: 15.33 ± 1.45 years; Cliff’s Delta = 1.00, p < 0.0001).

Correlations between the strength of the ISFS and clinical characteristics—including the duration of illness, age of onset, chlorpromazine equivalent, and symptomatology—were assessed separately for the COS and EOS groups due to above mentioned group differences in clinical characteristics and the use of different clinical interviews (SAPS/SANS versus PANSS). The mean strength of the ISFS was extracted from a central-parietal cluster of interest (C3, Cz, C4, P3, Pz, P4), where the presence and strength of the ISFS were most reduced in the patient groups. In the COS group, the strength of the ISFS showed a negative trend-level association with positive symptomatology (SAPS: rho = -0.47, p = 0.06). No other clinical characteristic was significantly correlated with the strength of the ISFS in either the COS group (age of onset: rho = 0.17, p = 0.51; duration of illness: rho = -0.33, p = 0.19; antipsychotic equivalent: rho = -0.07, p = 0.80; SANS: rho = -0.25, p = 0.34) or the EOS group (age of onset: rho = 0.19, p = 0.57; duration of illness: rho = 0.07, p = 0.84; antipsychotic equivalent: rho = 0.28, p = 0.41; PANSS Negative Score: rho = 0.05, p = 0.88; PANSS Positive Score: rho = 0.06, p = 0.86).

### Control analyses ensuring comparability of sleep data

To ensure that differences observed in all-night sleep structure parameters between the COS and HC-COS groups (reported in **3.1**) were not associated with the strength of the ISFS, the measures were correlated. Specifically, total sleep time, percentage of wake after sleep onset, and percentage of N1 sleep in the COS group were correlated with the mean strength of the ISFS from a central-parietal cluster of interest (C3, Cz, C4, P3, Pz, P4), where the presence and strength of the ISFS were most reduced in the patient groups. None of the sleep structure parameters (total sleep time: rho = 0.12, p = 0.65; wake after sleep onset: rho = -0.09, p = 0.73; N1: rho = 0.18, p = 0.48) showed a significant relationship with the mean strength of the ISFS.

To ensure comparability of N2 sleep bout data between the COS and HC-COS groups, the ISFS and sleep spindle analyses were repeated on an adjusted subset of data. Specifically, in the HC-COS group, the last 100 seconds were removed from bouts lasting longer than 400 seconds. This adjustment eliminated significant differences in N2 sleep bout data between the COS and HC-COS groups (**Supplementary Table 1**). The ISFS and sleep spindle density analyses were repeated where the COS and HC-COS groups had a comparable amount of data. Results showed similar patterns as those presented in **Figure 1** and **Figure 3** (**Supplementary Figure 1)**.

## Discussion

This study found that the infraslow fluctuation of sigma power (ISFS) is observable in young individuals with childhood-onset schizophrenia (COS) and early-onset schizophrenia (EOS). However, features of the ISFS, namely its presence and strength, are reduced in central-parietal regions in schizophrenia compared to healthy controls, suggesting less clustering of sleep spindles on an infraslow time scale. These alterations in the ISFS were observed in both the COS and EOS groups, with no significant differences between the two clinical groups despite differences in sleep spindle density and clinical characteristics.

Young healthy controls show a high proportion of electrodes where the ISFS is present, exhibiting a pronounced central-parietal topographical pattern. This aligns with recent findings showing that the ISFS is present from childhood to young adulthood, with its strength peaking in central-parietal regions (Dimitriades et al., 2024; Lecci et al., 2017). While the ISFS is observable in the COS and EOS groups, its presence and strength are notably reduced in these central-parietal regions compared to healthy controls. This is the first characterization of the ISFS in patient groups, particularly in individuals with schizophrenia. Therefore, a direct comparison of our findings with human data is limited. Evidence from rodent studies (Hayat et al., 2020; Osorio-Forero et al., 2021), along with recent human findings using pupil size as a proxy for locus coeruleus activity (Carro-Domínguez et al., 2025), suggests that the ISFS arises from the locus coeruleus-noradrenaline system, which regulates arousal and thalamocortical activity. Several lines of evidence point to dysfunction in the locus coeruleus-noradrenaline system in the pathophysiology of schizophrenia, contributing to cognitive deficits and symptomatology (Bird et al., 1980; Crow et al., 1979; Farley et al., 1978; Mäki-Marttunen et al., 2020; Maletic et al., 2017; Pelegrino et al., 2023; Steinhauer & Hakerem, 1992; Yamamoto & Hornykiewicz, 2004). In this context, the reduction in features of the ISFS in schizophrenia may stem from dysregulation of the locus coeruleus-noradrenaline system. However, the relationship between the ISFS and the pathophysiological mechanisms of schizophrenia requires further validation across species.

Markedly, both the COS and EOS groups show reduced ISFS features compared to healthy controls, with no significant differences between the two clinical groups, despite significant differences in sleep spindle density and clinical characteristics. Sleep spindle density was only significantly reduced in the COS group when compared to healthy controls and the EOS group, whereas no statistical difference was found between the EOS group and healthy controls. This aligns with our previous findings that the COS group shows robust, global reductions in sleep spindles compared to healthy controls, while the EOS group shows circumscribed, smaller reductions (Dimitriades et al., 2023). The lack of a significant difference between the EOS group and controls is likely due to methodological differences between our previous analysis (Dimitriades et al., 2023) and the current one, including the use of a broader spindle frequency band and continuous N2 bout data. As speculated (Dimitriades et al., 2023), the difference in sleep spindle density between schizophrenia groups is likely related to the duration of illness, which has been negatively correlated with spindle deficits in a meta-analysis (Lai et al., 2022).

The absence of significant spindle deficits in the EOS group, along with no significant correlations between the strength of the ISFS and clinical characteristics (e.g., age of onset, duration of illness, chlorpromazine equivalent, and symptom severity), may suggest that the ISFS disruption emerges early in schizophrenia. The ISFS is driven by noradrenergic activity in the locus coeruleus, which modulates thalamic depolarization and in turn, spindle generation (Hayat et al., 2020; Osorio-Forero et al., 2021). As schizophrenia progresses, noradrenergic dysregulation may impair thalamocortical circuits, further reducing spindle density and suggesting that the ISFS impairment precedes sleep spindle deficits. Additionally, given that sleep spindle features and thalamocortical circuits undergo dynamic changes during childhood and adolescence (Alkonyi et al., 2011; Fair et al., 2010; Kwon et al., 2023; Purcell et al., 2017), longitudinal studies are needed to disentangle the effects of these processes on one another and establish causal relationships, particularly since the cross-sectional nature of this data limits our ability to make such conclusions.

As the ISFS and sleep spindles are both related to sigma activity, it was possible that a severe spindle deficit would render the ISFS undetectable. However, the current findings rather support the notion that while the ISFS and sleep spindles are interrelated, they capture distinct aspects of brain activity. Specifically, the ISFS reflects the temporal clustering of sleep spindles on an infraslow time scale. This distinction is supported by their differing topographical distributions— the presence and strength of the ISFS peak in central-parietal regions, while sleep spindle density is maximal in frontal regions in our population—and by our control analysis, which found no significant correlation between the strength of the ISFS and sleep spindle density at a central electrode. Moreover, reduced ISFS strength does not simply indicate fewer spindles, but rather suggests a reduced clustering of spindles on an infraslow time scale.

This alteration in the infraslow timing of sigma activity could potentially have functional consequences. Arousals and markers of memory reactivation during sleep occur during specific ISFS phases in both rodents and humans, potentially linked either to spindle activity itself or to the broader neurophysiological milieu created by alternating phases of high and low noradrenaline (Dimitriades et al., 2024; Lecci et al., 2017). Although not examined in this study, adult patients with chronic schizophrenia have been reported to exhibit an increased frequency of arousals and less effective procedural sleep-dependent memory reactivation (Bagautdinova et al., 2023; Genzel et al., 2015; Manoach et al., 2004; Seeck-Hirschner et al., 2010). Thus, a reduced ISFS in schizophrenia may lead to impaired coordination of brain states needed for optimal arousal regulation and memory reactivation during sleep.

There are several limitations to consider when evaluating the findings of this study. The relative rarity of access to clinical populations with schizophrenia in this age group presented a challenge which resulted in small to moderate group sizes. To enhance robustness and minimize over-interpretation of potential group differences in a population with high inter-individual variability, a two-to-one matching strategy was employed. However, the sample size still limits the application of analytical methods that test multiple associations or interactions between variables. Furthermore, functional aspects of the ISFS could not be explored due to methodological constraints, including differing online filters affecting the slow-wave range, which prevented capturing slow wave-sleep spindle coupling (a marker of memory reactivation), as well as the lack of electromyographic data for detecting arousals, and the absence of memory readouts. Lastly, although most patients were on antipsychotic medication, previous studies suggest no direct link between spindle density and antipsychotics (Manoach et al., 2014).

However, the potential relationship between the ISFS and antipsychotic medication, particularly specific psychotropic agents, could not be further assessed in this sample due to the sample size and medication heterogeneity. Larger studies including individuals at elevated risk for developing schizophrenia and individuals with adult-onset schizophrenia are needed to extend and replicate these findings. Additionally, alterations of the ISFS may extend beyond schizophrenia. Therefore, studies including individuals with other mental disorders might further test the specificity of distinct ISFS impairments or combined impairments of timing and the number of the sleep spindles. Mental diseases for which alterations of locus coeruleus mediated noradrenergic firing during wake have been found and contextualized in pathophysiological stress-response models such as Post-Traumatic Stress or Major Depression Disorders could be of particular interest (Goddard et al., 2010; Grueschow et al., 2021; Naegeli et al., 2018; O’Donnell et al., 2004).

In summary, the current findings reveal a reduction in features of the ISFS in central-parietal regions in both young patients with schizophrenia during the early stages and those in the chronic course of the disease. This opens avenues for future research into the mechanisms, function, and extent of the ISFS reduction in schizophrenia, which could inform pathomechanistic models underlying the disorder as well as future diagnostic processes and interventions. Current developments and strategies for novel neuromodulation techniques targeting sleep oscillations may need to address not only the quantity but also the timing of sleep spindles and other sleep oscillations. Additionally, the specificity of the ISFS reduction— whether or not accompanied by a sleep spindle deficit—requires further investigation across mental disorders with noradrenergic dysregulation.

## Contributions

Data Collection: SF, SK, FP, FW, VJ, CV, SL, AB, DD, AM, JR, LT

Methodology: MED, ES, JA, MG, RH

Formal Analysis: MED

Supervision: RH, MG

Writing - Original Draft: MED

Writing - Editing: MED, RH, MG

Writing - Reviewing: MED, ES, JA, SF, SK, FP, FW, VJ, CV, SL, AB, DD, AM, JR, LT, RH, MG

## Data and Materials Availability

All data supporting the findings of this study are securely stored at servers of the University Children’s Hospital. Access and availability will be provided upon a material transfer agreement and after approval by the local ethics committee of the Canton of Zurich.

## Acknowledgements

We would like to thank the participants for their time and data contributions, as well as their families for their support. We also appreciate the invaluable advice and support from everyone in Reto Huber’s lab and Miriam Gerstenberg’s research group throughout this project. This work was supported by the UZH Candoc Grant (to MED), Swiss National Science Foundation PCEFP1-181279 (to SK), 32003B_184943 (to LT), 320030_153387 (to RH), Interfaculty Research Cooperation: Decoding Sleep (to LT), and the Frutiger Foundation and the EMDO Foundation (to MG). This research was also supported in part by the Intramural Research Program of the National Institute of Mental Health (Annual Report Number ZIAMH002581, ClinicalTrials.gov identifier NCT00001198, protocol ID 84-M-0050).

