## Supplemental Table 1 and Figure 1 for "The Infraslow Fluctuation of Sigma Power During Sleep in Young Individuals with Schizophrenia"

### Supplementary Material

**Supplementary Table 1:** Comparable non-rapid eye movement sleep stage N2 bout data between the childhood-onset schizophrenia group and age- and sex-matched healthy controls

| **Parameter** | **Patient Group (N)** | **Mean (SD)** | **Healthy Controls (N)** | **Mean (SD)** | **Cliff's Delta** | **p** |
|---|---|---|---|---|---|---|
| <u>N2 Bout Sleep</u> | | | | | | |
| Bout Number | COS (17) | 13.71 (10.09) | HC-COS (34) | 12.71 (3.24) | 0.02 | 0.92 |
| Mean Bout Length (s) | COS (17) | 450.58 (86.42) | HC-COS (34) | 510.69 (124.07) | -0.30 | 0.08 |
| Mean Relative Location (%) | COS (17) | 52.23 (13.34) | HC-COS (34) | 54.03 (7.14) | -0.06 | 0.73 |
| Total Duration of Bouts (min) | COS (17) | 110.35 (91.19) | HC-COS (34) | 110.83 (43.70) | -0.08 | 0.65 |
| Percent of all N2 | COS (17) | 38.23 (23.40) | HC-COS (34) | 47.91 (15.07) | -0.23 | 0.18 |

**Supplementary Table 1:** COS: Childhood-Onset Schizophrenia; HC-COS: Healthy Controls for COS; SD: Standard Deviation; N2: Non-Rapid Eye Movement Sleep Stage N2; min: minutes; s: seconds; %: percent. Bout: N2 sleep data lasting at least 300 seconds without transitions to other vigilance states or artifacts. Bouts of N2 Sleep that lasted 400 seconds or longer in the HC-COS group were reduced by 100 seconds in order to make the amount of bout data comparable between the COS and HC-COS groups. Mean Relative Location: Mean of all bout midpoints divided by the total recording length, expressed as a percentage. Proportion of all N2: Total duration of bout data relative to total N2 sleep duration. Statistically significant p-values are shown in bold.

**Supplementary Figure 1:** Analyses of features of the infraslow fluctuation of sigma power and sleep spindle density in comparable non-rapid eye movement stage N2 sleep bout data

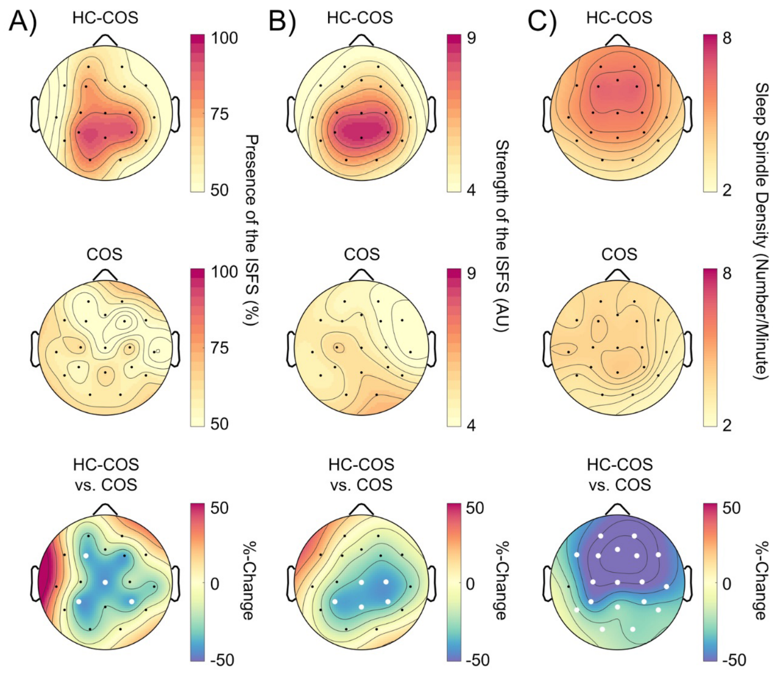


**Supplementary Figure 1**: Topographical distribution of the A) presence of the infraslow fluctuation of sigma power (ISFS), B) strength of the ISFS, and C) sleep spindle density between the Childhood-Onset Schizophrenia (COS; middle row) and their respective healthy control group (HC-COS; top row) with comparable Non-Rapid Eye Movement Stage N2 Sleep Bout Data. Specifically, in the HC-COS group, the last 100 seconds were removed from bouts lasting longer than 400 seconds. This adjustment eliminated significant differences in N2 bout data between the COS and HC-COS groups. Presence of the ISFS is expressed as the percentage of individuals in each group exhibiting the ISFS at a given electrode. Group differences in the presence of the ISFS were assessed using a z-test for proportions (electrodes showing significant group differences are marked with a white dot) and presented as the percent change between groups (bottom left). The strength of the ISFS and sleep spindle density were compared using an unpaired t-test (electrodes showing significant group differences are marked with a white dot) and are presented as the percent change between groups (bottom middle and right, respectively).
